# Mito-SiPE: A sequence-independent, PCR-free mitochondrial DNA enrichment method for ultra-deep sequencing that minimises amplification and alignment artefacts for the analysis of mitochondrial heteroplasmy/variation

**DOI:** 10.1101/2022.10.15.512094

**Authors:** Darren J Walsh, David J Bernard, Faith Pangilinan, Madison Esposito, Denise Harold, Anne Parle-McDermott, Lawrence C Brody

**Affiliations:** Gene and Environment Interaction Section, National Human Genome Research Institute, NIH, Bethesda, MD, USA; School of Biotechnology, Dublin City University, Dublin, Ireland

## Abstract

**Background:** Deep sequencing is often used to measure somatic variation in the mitochondrial genome. Selective enrichment methods, such as PCR amplification or probe hybridization/capture are commonly used. These methods can introduce bias and are prone to contamination by nuclear-mitochondrial sequences (NUMTs); elements that can introduce artefacts into analyses such as an assessment of mitochondrial heteroplasmy.

**Results:** Here, we demonstrate a method to obtain ultra-deep (>80,000X) sequencing coverage of the mitochondrial genome by selectively purifying the intact organelle itself using differential centrifugation and alkaline lysis. We applied this approach to seven different mouse tissues. Isolation of mitochondria yields a preparation of highly enriched mtDNA. We compared this method to the commonly used PCR-based method. Mito-SiPE avoids false-heteroplasmy calls that occur when long-range PCR amplification is used for mtDNA enrichment.

**Discussion:** We have described a modified version of a long-established protocol for purifying mtDNA and have quantified the increased level of mitochondrial DNA post-enrichment in 7 different mouse tissues. This method will enable researchers to identify changes in low-frequency heteroplasmy without introducing PCR biases or NUMT contamination that are falsely identified as heteroplasmy when long-range PCR is used.

## Background

Decades of research have established a link between mitochondrial DNA variation and human health. Recent advances in DNA sequencing technologies have led to an increased ability to interrogate the mitochondrial genome for low-frequency mutations associated with various disease states. Mitochondrial DNA mutations have been associated with ageing^1^ and a myriad of disease phenotypes^2^. Disorders caused by inherited and acquired mitochondrial DNA variants affect ~1 in 4,300 of the population^3^. These variants were initially thought to solely originate from matrilineal inheritance of mitochondrial DNA molecules, however more recent studies have shown that somatic mutations also occur in mtDNA over time and in a tissue-specific manner^4–6^. There are hundreds-to-thousands of mitochondrial DNA molecules in every human cell. This number is dependent on the tissue, cell type and energy state of the mitochondria^7^. The fluctuating, multi-copy nature of mitochondrial DNA means that mutations can be present at any frequency within a cell, unlike the diploid nuclear genome. The presence and frequency of mitochondrial DNA mutations is referred to as mitochondrial heteroplasmy. While there appears to be a threshold frequency for the impact of some heteroplasmic variants on human health in relation to some conditions; the specific variants and their required pathogenic frequency has yet to be established for acquired somatic mitochondrial mutations^8–12^.

Mitochondrial heteroplasmy has been increasingly investigated as a contributor to human disease. To date, it has been linked to various diseases including cardiomyopathy, hypertension, epilepsy, Parkinsons disease and optic neuropathy^8–12^. Elevated heteroplasmy levels have also been linked to tumor aggressiveness and poor cancer prognosis^13,14^. These studies demonstrate that high resolution mitochondrial heteroplasmy analyses could be used to identify molecular mechanisms that drive some disease states. Additionally, evidence suggests that mitochondrial DNA heteroplasmy could be used for diagnosis/prognosis of particular conditions and even perhaps as a therapeutic target^15^. Given these findings, it is important that mitochondrial heteroplasmy, particularly low frequency heteroplasmy, can be identified and quantified as a part of disease-related studies. Such investigations require high to ultra-high sequencing coverage (>1000X - >10,000X) of the mitochondrial genome in order to reliably quantitate low frequency heteroplasmy with a high degree of sensitivity and specificity. Currently, this is typically achieved by using probe hybridisation/capture or long-range polymerase chain reaction (PCR) to enrich mitochondrial DNA.

Probe hybridisation/capture^16^ uses complementary probes that bind mitochondrial sequences to separate mtDNA from nuclear DNA. Another approach is long-range PCR^17^ which amplifies the entire mitochondrial genome, typically in two, overlapping fragments. Both probe hybridization and long-range PCR amplification require complementary binding of probes/primers to enrich mtDNA from whole DNA extracts. Widespread use of these methodologies in many heteroplasmy studies is due to their ease, amenability to high-throughput processes and efficacy of producing mtDNA appropriate for ultra-deep sequencing^4,5,18,19^. These sequence-dependent methods are imperfect. Probes and primers designed to match reference alleles may select against rare heteroplasmic variants that are of interest. Additionally, PCR amplification is known to introduce errors that may appear as false positive heteroplasmic variants. Arguably, the most problematic issue for these techniques is the contamination of nuclear-mitochondrial elements (NUMTs)^20^.

NUMTs are nuclear sequences that share high levels of sequence identity with mtDNA. They arose from the somatic translocation of mitochondrial DNA into the nuclear genome. The number, size and sequence of NUMTs varies within species^21^, including between human populations and indivduals^22^. The entire mitochondrial genome is represented in the human nuclear genome^22^. As a result, it is extremely difficult to design primers or hybridization probes that will selectively enrich mitochondrial DNA without also enriching NUMT sequences. Multiple studies have found that NUMT contamination was present in mtDNA sequencing data that used either probe hybridisation or PCR amplification to enrich mtDNA^23–26^. Most notably, NUMT contamination is thought to explain the apparent paternal inheritance of mitochondrial DNA in humans that was reported in PNAS in 2018^27–30^. The difficulty posed by deciphering NUMT contamination from true mitochondrial DNA may require the use of a sequence-independent capture method of mitochondrial enrichment.

In this study we have adapted a previously described^31^ sequence-independent and PCR-free technique which relies on differential centrifugation and alkaline lysis to separate mitochondria from other tissue/cellular debris. We provide evidence that this method can be effectively used to isolate mitochondrial DNA from different tissue types for subsequent mtDNA sequencing. We achieved ultra-deep coverage of the mitochondrial genome when this approach was combined with an appropriate Next-Generation Sequencing (NGS) data pipeline. To demonstrate the proof-of-concept, we employed *Polg* mutator mice as positive controls to compare this methodology with long-range PCR amplification enrichment of mtDNA. These mice lack the ability to ‘proof-read’ their mitochondrial DNA and as a result, gather single nucleotide mutations at a much higher frequency than their wild-type counterparts^32^. We propose that this method can be applied to a range of species to allow researchers to reliably investigate mitochondrial heteroplasmy and expand our current knowledge of the contribution of somatic mitochondrial mutations to human ageing and disease. This methodology negates the impact of NUMT contamination and PCR error introduction when assessing heteroplasmy and is therefore more sensitive and accurate than long-range PCR amplification. We refer to this method as Mito-SiPE.

## RESULTS

The Mito-SiPE technique for enriching mtDNA for sequencing is rooted in classic cell biology and biochemistry methods for subcellular fractionation. It utilizes a combination of differential centrifugation and alkaline lysis to separate the mitochondria from nuclear and cytoplasmic cellular components. The resulting preparation is then used to purify mitochondrial DNA with minimal contamination from nuclear DNA (Fig. 1a). To test this method across a range of inputs, we used seven different mouse tissues; brain, heart, lung, kidney, liver, spleen and muscle. Mitochondrial copy number was assessed *via* quantitative polymerase chain reaction (qPCR) in enriched samples from the seven tissues and compared to unenriched, whole DNA extracts (Fig. 1b). qPCR was used to calculate the ratio of mitochondrial DNA to nuclear DNA, which provides an estimate of the average mtDNA copy number per cell. A significant increase (100-1,000 fold, P < 0.0005) in mitochondrial DNA copy number was observed in samples that underwent enrichment. This effect was present across the 7 tissues of interest (Fig. 1c).

**Figure 1.**
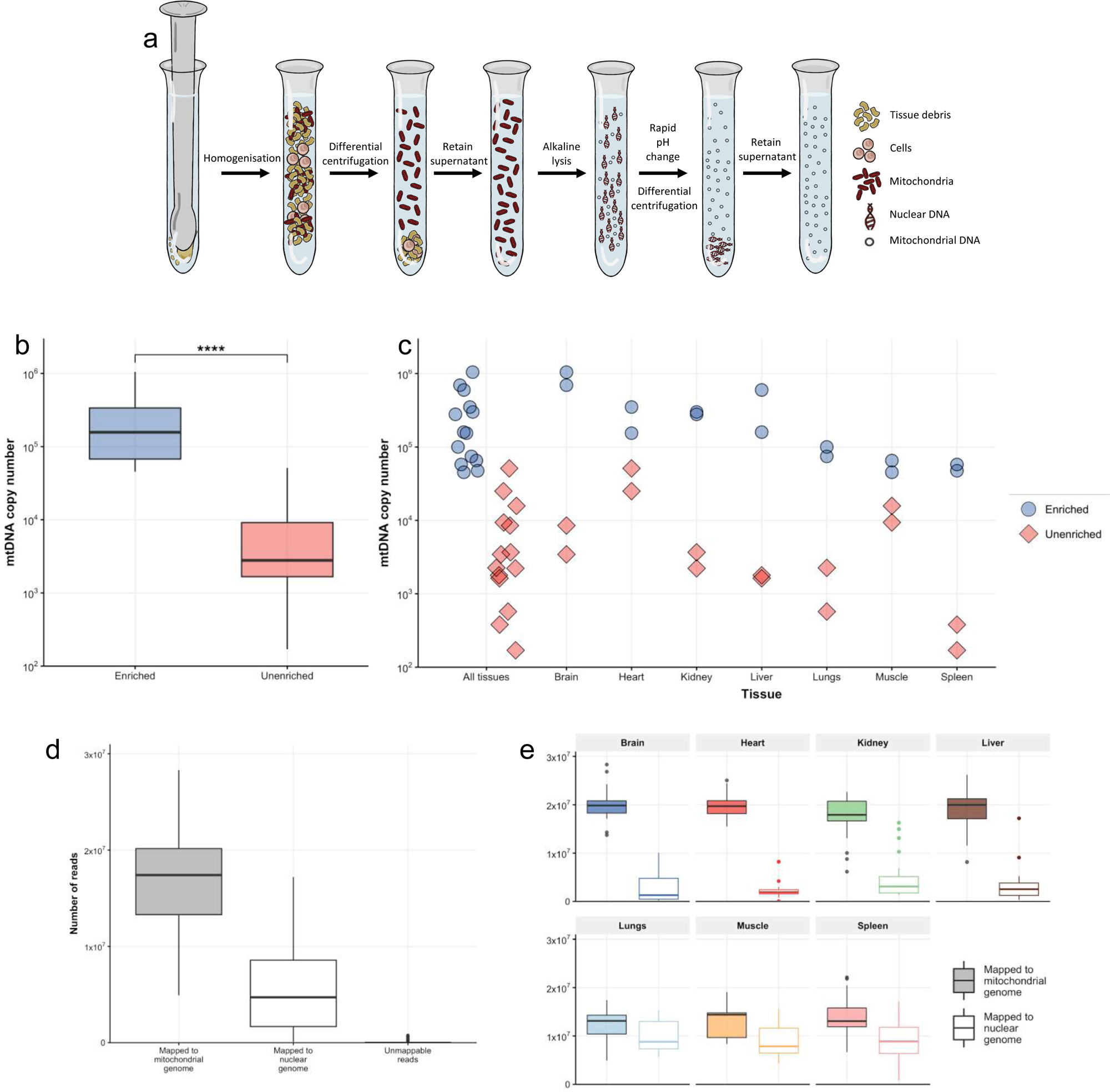
PCR-free enrichment of mitochondrial DNA using differential centrifugation and alkaline lysis. **a**, Overview of sequence-independent and PCR-free mitochondrial DNA enrichment workflow. Homogenisation and differential centrifugation are used to enrich mitochondria. Alkaline lysis is then used to isolate mitochondrial DNA from any remaining nuclear DNA. **b**, Relative quantification of mtDNA copy number from seven different mouse tissues that underwent enrichment. Mitochondrial DNA to nuclear DNA ratio (mtDNA:nuDNA) was assessed *via* qPCR. P < 0.0005 Wilcoxon signed rank test. Log scaled. **c**, The mtDNA:nuDNA ratio across seven different tissues which underwent enrichment. (n=2) for each tissue (brain, heart, lungs, liver, kidney, spleen and muscle). **d**, The distribution of nuclear, mitochondrial and unmappable reads generated in each sample displayed as boxplots (n=163). **e**, The distribution of total and mapped reads across seven mouse tissues after mtDNA enrichment; Brain (n=26), Heart (n=26), Kidney (n=25), Liver (n=21), Lungs (n=26), Muscle (n=12) and Spleen (n=27).

### Comparison of two alignment strategies to assess nuclear contamination

The performance of this technique to produce pure, high-quality mtDNA for next-generation DNA sequencing was assessed by applying the method to 163 samples across seven different mouse tissues. These samples were then sequenced across four lanes of the Illumina NovaSeq platform. After quality control, alignment and removal of duplicates, the number of mitochondrial, nuclear and unmapped reads were assessed for each sample (Fig. 1d, Table 1). Of the mapped reads, an average of 75% ±20% were mapped to the mitochondrial genome and 25% ±20% to the nuclear genome. Unmappable reads made up 0.26±0.63% of the total reads (Table 2). The level of nuclear contamination in samples originating from brain, heart, kidney and liver was extremely low with higher levels found in lung, muscle and spleen (Fig. 1e).

**Table 1.** The number of mitochondrial, nuclear and unmapped reads per tissue.

| ***Tissue*** | **Mitochondrial (sd)** | **Nuclear (sd)** | **Unmapped (sd)** |
| --- | --- | --- | --- |
| *Brain* | 20.027 (3.285) | 2.575 (2.729) | 0.073 (0.178) |
| *Heart* | 19.636 (2.166) | 2.188 (1.481) | 0.042 (0.099) |
| *Kidney* | 17.428 (4.224) | 4.783 (4.344) | 0.012 (0.021) |
| *Liver* | 18.931 (4.071) | 3.291 (3.774) | 0.081 (0.171) |
| *Lungs* | 12.206 (2.938) | 9.860 (3.137) | 0.057 (0.134) |
| *Muscle* | 13.146 (3.467) | 9.093 (3.589) | 0.156 (0.252) |
| *Spleen* | 13.813 (3.827) | 9.301 (4.368) | 0.037 (0.072) |
| ***All Tissues*** | **16.641 (4.586)** | **5.700 (4.660)** | **0.057 (0.139)** |

**Table 2.**
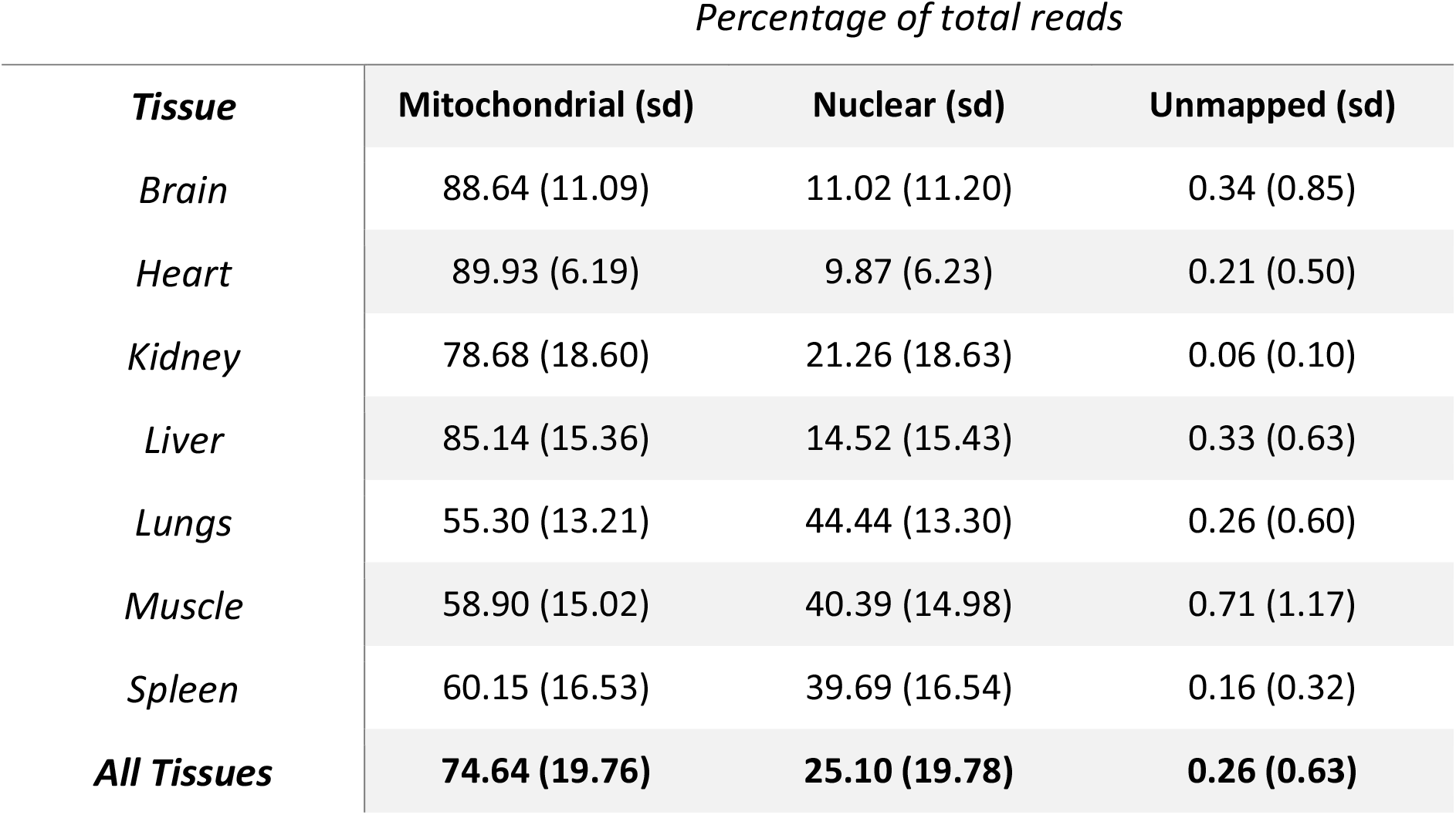
The percentage of reads aligned to the mitochondrial genome, nuclear genome and unmappable reads.

Two alignment strategies were used to assess the coverage of the mitochondrial genome and the distribution of nuclear contamination (Fig. 2a). The first method aligned filtered reads to the whole genome and then isolated mitochondrial-aligned reads. The second method aligned all the reads to the mitochondrial genome first, then mapped any remaining unaligned reads to the nuclear genome. Coverage across the mitochondrial genome and distribution of nuclear contamination were assessed after duplicate removal. Average coverage across the mitochondrial genome exceeded 50,000X at each base pair and was dependent on tissue of origin (Fig 2b). Lung, muscle and spleen had lower levels of coverage due to the higher levels of reads mapped to nuclear DNA in these samples. Mapping to the whole genome first caused a loss in coverage between nucleotide positions 7,500-11,000 (Fig. 2b, top). This dip in coverage was not observed when reads were first aligned to the mitochondrial genome using the second alignment strategy (Fig. 2b, bottom). The latter method did not lead to an overall increase in sequencing coverage across the rest of the mitochondrial genome, indicating that nuclear reads were not misaligned to the mitochondrial genome when this method was used.

**Figure 2.**
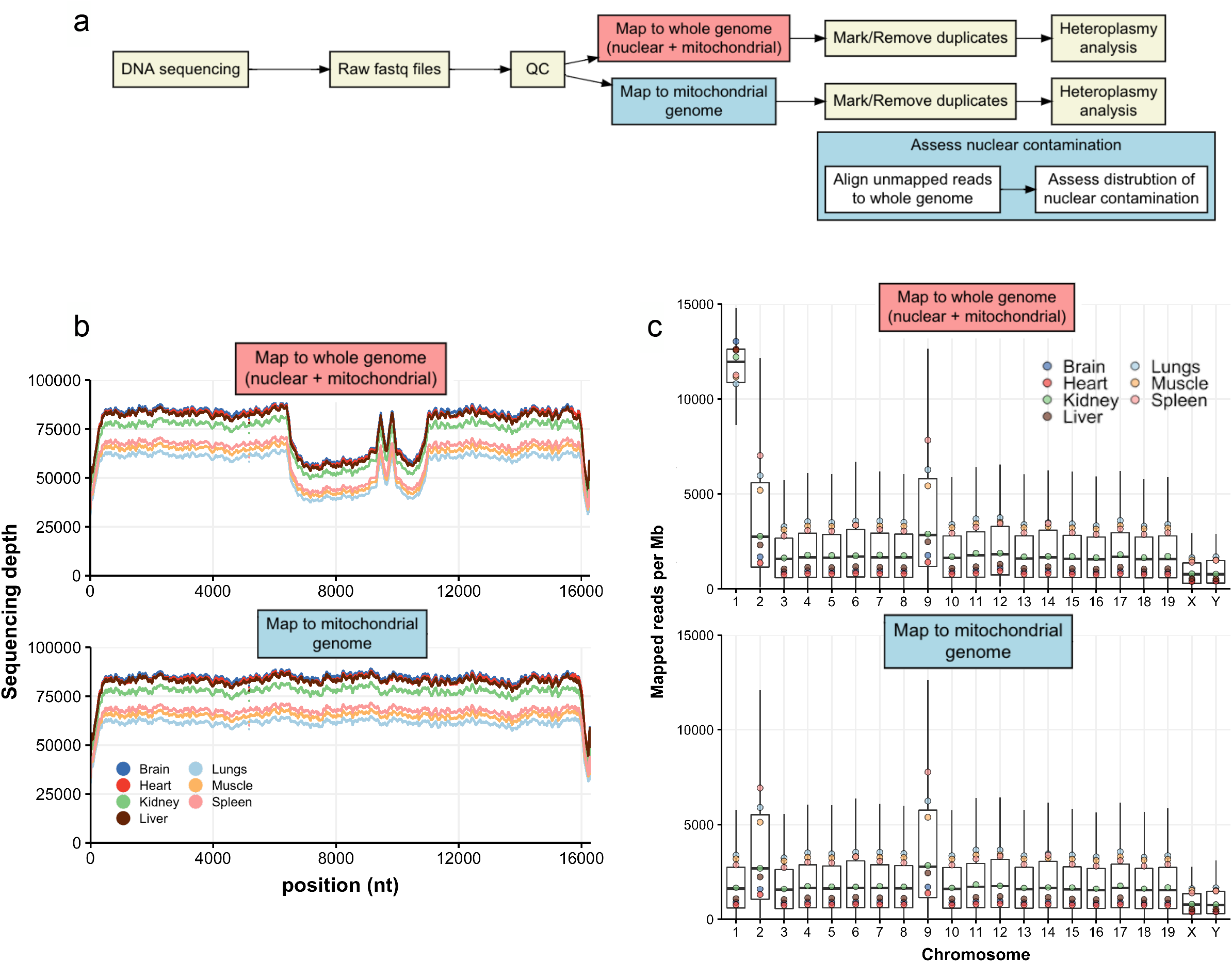
Alignment of sequencing data to whole-genome reference leads to misalignment of mitochondrial reads. Contaminating nuclear reads are randomly distributed across the whole genome. **a**, Overview of two methods for mapping sequencing data; *via* whole genome mapping or mapping exclusively to the mitochondrial genome followed by mapping of unaligned reads to the nuclear genome. **b**, Top, sequencing coverage across the mitochondrial genome of seven mouse tissues when reads are mapped to the whole reference genome. A ‘hole’ in coverage is observed between nucleotide positions 7,500-11,000. Bottom, sequencing coverage across the mitochondrial genome of seven mouse tissues when reads are mapped exclusively to the mitochondrial reference genome. **c**, Top, the box plots represent the distribution of reads mapped per megabase of DNA to each chromosome when reads are aligned to the whole reference genome. Bottom, the box plots represent the distribution of reads mapped per megabase of DNA to each chromosome when reads are aligned exclusively to the mitochondrial genome. The points represent the average reads per Mb mapped for each tissue.

As this mtDNA enrichment method is sequence-independent, any reads originating from nuclear DNA should have been randomly distributed throughout the whole genome. To test this hypothesis, the distribution of nuclear contamination was assessed using both alignment strategies (Fig 2c). When reads were aligned to the whole genome, nuclear contamination appeared to be evenly distributed across the nuclear genome as expected, except for chromosome 1 (Fig 2c, top). Aligning the reads to the mitochondrial genome first did not produce this same anomaly (Fig 2c, bottom and Supplementary Fig. 1). This is due to an NUMT found on chromosome 1 of the reference genome that shares 100% sequence identity of its homologous sequence in the mitochondrial genome. As a result, the alignment tool (bwa) mapped ~50% of reads originating from this region to the mitochondrial genome and ~50% incorrectly, to the homologous NUMT on chromosome 1 when reads were mapped to the whole genome. This artefact was eliminated when reads were mapped to the mitochondrial genome first. Chromosomes 2 and 9 also had slightly elevated levels of mapped reads when compared to the rest of the genome. This effect was produced by both alignment strategies. However, further investigation showed that this was caused by the alignment of highly repetitive reads from across the genome to the same loci on chromosomes 2 and 9 (Supplementary Fig 2,3 and Supplementary Fig 4,5, respectively). Mapping to the mitochondrial genome first appeared to be more effective at mapping true mitochondrial reads correctly, without increasing levels of spurious alignments. This alignment methodology was used to assess heteroplasmy levels in the tissues of *Polg*^D257A/D257A^ and *Polg*^wt/wt^ mice to demonstrate the proof-of-concept of Mito-SiPE.

### Sequence coverage and heteroplasmy in samples prepared with Mito-SiPE compared to lrPCR

Mito-SiPE and long-range PCR (lrPCR) were applied to the Polg mice. There was an average of 7.15 × 10^7^ reads produced per sample across all studied groups (Table 3; n=48). A higher average number of reads was observed in the lrPCR amplified samples (7.22 and 7.19 × 10^7^, *Polg*^D257A/D257A^ (n=12) and *Polg*^wt/wt^ (n=12), respectively) compared to the Mito-SiPE samples (7.15 and 7.02 × 10^7^ *Polg*^D257A/D257A^ and *Polg*^wt/wt^, respectively). The proportion of reads mapped to the mitochondrial genome was also higher in the lrPCR amplification samples than the Mito-SiPE samples (Table 3). The average coverage across the mitochondrial genome was higher in the lrPCR amplified samples (average depth 137,000X compared to the Mito-SiPE average of 123,000X). However, these samples had a loss in coverage towards the end of the two overlapping fragments (Fig 3a, left). The Mito-SiPE samples, broadly, had uniform coverage across the entire mitochondrial genome compared to the lrPCR samples. *Polg*^D257A/D257A^ samples displayed minor region-specific fluctuations in coverage (Fig 3a, right). This effect was not observed in the samples amplified with lrPCR (Fig 3a, middle).

**Table 3.** The average total number of reads, mapped reads and sequencing depth for each methodology and genotype.

| Assay type | Genotype | Total reads |  | Mapped reads |  | Coverage |  |
| --- | --- | --- | --- | --- | --- | --- | --- |
|  |  | Average (10<sup>7</sup>) | SD | Average (10<sup>7</sup>) | SD | Average (10<sup>5</sup>) | SD |
| IrPCR amplification | *Polg*<sup>D257A</sup> | 7.22 | 0.561 | 7.19 | 4.50 | 1.39 | 0.417 |
| mtDNA prep | *Polg*<sup>D257A</sup> | 7.15 | 0.848 | 3.98 | 1.86 | 1.22 | 0.245 |
| IrPCR amplification | *Polg*<sup>wt</sup> | 7.19 | 0.687 | 7.08 | 0.693 | 1.36 | 0.411 |
| mtDNA prep | *Polg*<sup>wt</sup> | 7.02 | 0.687 | 4.53 | 1.79 | 1.25 | 0.126 |

**Figure 3.**
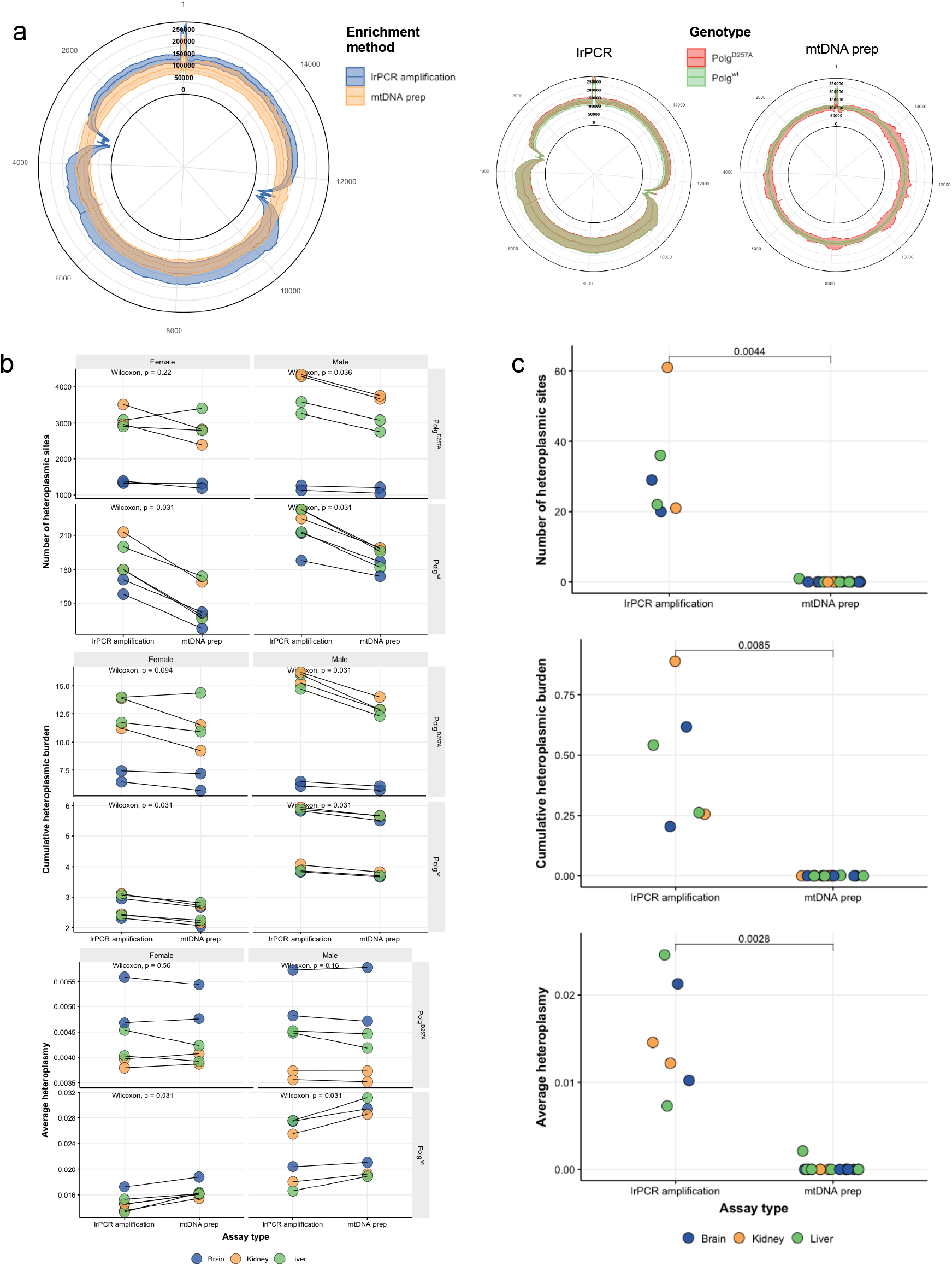
Mitochondrial DNA preparations outperform long-range PCR amplification and reduce the impact of PCR errors and NUMT contamination on mitochondrial heteroplasmy. **a**, Left, read coverage across the mitochondrial genome after sequencing and alignment for each enrichment method. The average sequencing depth was higher in the lrPCR (blue) samples. There was a significant reduction of coverage towards the end of each amplicon fragment in the lrPCR samples. Right, differences in the average sequencing depth between Polg^wt/wt^ (green) and Polg^D257A/D257A^ (red) using mtDNA prep (left) and lrPCR (right) methodologies. The sequencing depth of Polg^D257A/D257A^ tissues enriched using the mtDNA prep method showed a region-specific variation in coverage, however the average coverage remained comparable between both genotypes. This region-specific pattern was not observed in samples enriched using lrPCR. The standard deviations were larger in lrPCR enriched samples, with one fragment showing larger differences than the other. This was likely due to PCR efficiency or variations due to attempts to mix both fragments in equimolar ratios. **b**, top, the number of heteroplasmic sites (alternative allele frequency ≥ 0.2 %) that were identified in each sample. *Polg*^*D257A/D257A*^ tissues had more heteroplasmic sites than *Polg*^wt/wt^. There were significantly less heteroplasmic sites observed in mtDNA prep samples of *Polg*^wt/wt^ males and females, and in *Polg*^*D257A/D257A*^ males than in lrPCR enriched samples. This effect was not significant in the *Polg*^*D257A/D257A*^ females. Middle, The cumulative heteroplasmic burden identified across all samples using both methodologies. Cumulative heteroplasmic burden displayed a similar pattern to the number of heteroplasmic sites; significantly lower levels were detected in *Polg*^wt/wt^ males and females and in *Polg*^*D257A/D257A*^ males. The difference was not significant in *Polg*^*D257A/D257A*^ females. Bottom, The average alternative allele frequency (average heteroplasmy) observed across all samples using both enrichment methodologies. Average alternative allele frequency displayed a different pattern of results compared to the previous two heteroplasmy metrics. *Polg*^*D257A/D257A*^ tissues had lower mean alternative allele frequencies than *Polg*^*wt*^. Significantly higher mean alternative allele frequencies were observed in lrPCR tissues from both male and female *Polg*^*D257A/D257A*^ mice. Conversely, *Polg*^wt/wt^ samples enriched using the mtDNA prep methodology had higher mean alternative allele frequency than those enriched via lrPCR. Tissues are highlighted by colour, each line represents the same tissue that was enriched using both methodologies. Statistical comparisons between lrPCR and mtDNA prep were performed using a Wilcoxon signed-rank test. There were 24 lrPCR samples and 24 mtDNA prep samples (*Polg*^wt/wt^ n=24, *Polg*^*D257A/D257A*^ n=24). Two samples for each tissue, sex and genotype were analysed. **c**, Heteroplasmy levels of C57BL6 wild-type mice using both lrPCR and mtDNA prep enrichment methods. Mitochondrial DNA from C57BL6 tissues that were enriched using lrPCR had substantially higher levels of heteroplasmy across the three assessed metrics; top, number of heteroplasmic sites, middle, cumulative heteroplasmic burden and bottom, average heteroplasmy. Unlike the Polg mutator mouse comparisons, these enrichments were not performed on the same tissues however the mice were the same sex and age at sacrifice. Statistical comparisons between lrPCR and mtDNA prep were performed using a paired Student’s T-test. There were six samples in the lrPCR group and twelve samples in the mtDNA prep group.

Three metrics were measured to assess heteroplasmy levels in *Polg*^D257A/D257A^ and *Polg*^wt/wt^ tissues: the number of heteroplasmic sites, the average alternative allele frequency (heteroplasmy frequency) and cumulative heteroplasmic burden. These three metrics are not independent and have previously been reported in the literature^14,33–37^. The number of heteroplasmic sites is the number of nucleotide positions at which an alternative allele was identified above the threshold frequency (0.2%) in a sample. Alternative allelic calls caused by sequencing error are present below this frequency and are indistinguishable from low-frequency heteroplasmy. Average heteroplasmy is the mean frequency of all variants observed in a sample above the threshold frequency. Finally, cumulative heteroplasmic burden is the sum of all variant frequencies that were identified above the threshold frequency in a sample.

More heteroplasmic sites were observed in the *Polg*^*D257A/D257A*^ mice than wild-type, in brain, liver and kidney (Fig 3b, top). No difference was observed between *Polg*^*D257A/D257A*^ males and females, however there was a significant difference between sexes of *Polg*^*wt/wt*^ (Fig 3b, top, left panels vs right panels). There were significantly fewer heteroplasmic sites found in Mito-SiPE preps of both male and female *Polg*^*wt/wt*^ as well as *Polg*^*D257A/D257A*^ males compared to lrPCR enrichment. This effect was not statistically significant in female *Polg*^*D257A/D257A*^ tissues. The difference between enrichment assays was larger in *Polg*^*D257A/D257A*^ samples than in *Polg*^*wt/wt*^ (Supplementary Fig 6). When examining cumulative heteroplasmic burden, similar overall results were observed (Fig 3b, middle). Lower burden was detected in Mito-SiPE prep enriched samples compared to lrPCR in the same 3 of 4 comparisons. Average heteroplasmy, however, displayed contrasting results (Figure 3b, bottom). *Polg*^*D257A/D257A*^ mice had lower average heteroplasmy levels than *Polg*^*wt/wt*^ mice. In *Polg*^*D257A/D257A*^ mice, the mean alternative allele frequency was significantly lower in Mito-SiPE preps than in lrPCR enrichment samples. *Polg*^*wt/wt*^ mice appeared to have higher average heteroplasmy levels, which were elevated in mtDNA preps compared to lrPCR amplification enrichment. Although this is a surprising result, it is caused by the many low-frequency heteroplasmic variants that are present in *Polg*^*D257A/D257A*^ tissues.

### mtDNA mutation rate comparison in *Polg*^*D257A/D257A*^ mice compared to C57BL6/J mice

*Polg*^*wt/wt*^ mice have a baseline of mitochondrial DNA mutation much higher than that of true wild-type C57BL6 mice (169 ± 29.4 and 0.06 ± 0.24 heteroplasmic sites, respectively). This is because heterozygous female breeders have an intermediate phenotype. Long-range PCR amplification enrichment was performed on total DNA extracted from brain, kidney and liver of two wild-type C57BL6/J males. These enrichments were compared to mtDNA preparation of 6 control C57BL6/J males to investigate the effect of methodology on wild-type mice without a *Polg* background as they have a higher baseline of heteroplasmy than wild-type. These mice were age-matched. There was a significant and substantial increase in the apparent number of heteroplasmic sites, cumulative heteroplasmic burden and average heteroplasmy in the lrPCR samples compared to samples that underwent mtDNA preparation (Figure 3c).

### Mito-SiPE analysis across tissues of *Polg*^*wt/wt*^ and *Polg*^*D257A/D257A*^ mice

Mito-SiPE was performed on 7 tissues of *Polg*^*wt/wt*^ and *Polg*^*D257A/D257A*^ mice. Mutant mice displayed significantly higher levels of mitochondrial DNA mutation than wild-type across all tissues (Supplementary Fig 7). Kidney, liver, colon, heart and lung had a similar number of heteroplasmic sites in *Polg*^*D257A/D257A*^ mice with higher levels observed in spleen and lower levels found in the brain (Supplementary Fig 8a). Colon and spleen had higher levels of average heteroplasmy than the other five tissues in *Polg*^*D257A/D257A*^ mice (Supplementary Fig 8b). Cumulative heteroplasmic burden was higher in *Polg*^*D257A/D257A*^ colon and spleen tissues and lower in brain than in heart, kidney, liver and lung (Supplementary Fig 8c). There was no significant difference between the tissues of *Polg*^*wt/wt*^ mice across any of the three heteroplasmy metrics that were assessed.

## Discussion

In this study we have demonstrated that a sequence-independent technique for mitochondrial DNA enrichment (that we refer to as Mito-SiPE) is highly effective and can produce ultra-deep sequencing coverage required for heteroplasmy analysis. We conclude that this methodology works most effectively with brain, heart, liver and kidney samples, however sufficient results can also be obtained in samples originating from lung, muscle and spleen. The cause of the disparity between tissues is unknown, however, we hypothesise that it may be related to the amount of starting material, the mechanical properties of the tissues and mitochondrial copy number before enrichment. There are three main advantages of this method over the current standard: 1) It does not require complementary binding of reference probes/primers to mitochondrial DNA, 2) PCR amplification is not required and therefore generates no polymerase errors during enrichment and 3) Any nuclear contamination present is randomly distributed across the nuclear genome and therefore does not result in NUMT enrichment leading to false interpretation of any resulting data.

The sequencing depth achieved using this methodology exceeds that of many heteroplasmy studies to date, even in the tissues where enrichment was less effective^5,19,38,39^. High coverage of the mitochondrial genome allows for a more sensitive assessment of low frequency mutations and changes in heteroplasmy frequency. The number of reads (average 22.4 million per sample, Table 1) produced through sequencing in this study could be reduced by up-to 5-fold and would still be sufficient to achieve coverage levels in line with previous studies (typically less than 10,000X). This would reduce the sequencing costs, increase throughput and enable the study of mitochondrial heteroplasmy across many samples by taking advantage of high-throughput sequencing. Our results also suggest that sufficient sequencing coverage could be achieved even with lower-capacity sequencing platforms.

An alternative bioinformatics pipeline is required when this technique is utilized. Typically, it is advised to align all sequencing data to the whole reference genome before isolating mitochondrial reads to avoid spurious alignment of nuclear reads to the mitochondrial genome^40^. However, due to the lack of sequence-specific enrichment when using this technique, we demonstrate that a different approach is optimal. When reads are aligned to the reference genome first, mitochondrial reads are incorrectly aligned to homologous regions in the nuclear genome, most prominently, chromosome 1. This effect in this study, it should be noted, is specific to the mouse reference genome; however, future studies may assess whether the same effect is observed in human alignments or indeed in other species. As we demonstrate here, mapping reads to the mitochondrial genome first did not appear to lead to an increase in spurious alignment of nuclear reads to the mitochondrial genome and produced uniform sequencing coverage.

Nuclear-mitochondrial sequences have been identified as an important source of artefacts in heteroplasmy analysis. By employing Mito-SiPE, we find that any nuclear contamination present in the resultant sequence data is randomly distributed across the nuclear genome. It is pertinent to note, however, that although NUMT contamination is not enriched in this study, it does not mean that NUMT contamination is entirely absent. This is an important point as the number and size of all NUMTs have not been fully elucidated for most species and varies within species, and even between individuals. The effect of NUMT contamination when using this technique will be minimised in the tissues that contain less nuclear contamination e.g., brain, heart, kidney and liver.

Mitochondrial DNA heteroplasmy has been assessed in different ways across a number of studies. Number of heteroplasmic sites, average alternative allele frequency and cumulative heteroplasmic burden are all metrics that have been considered^14,33–37^. The number of heteroplasmic sites and cumulative heteroplasmic burden was higher in *Polg*^D257A/D257A^ mouse tissues than in that of *Polg*^wt/wt^, using both lrPCR amplification and Mito-SiPE preparation methods. These findings are in line with previous studies, however the number of mutations detected appear to be higher in the data presented here. This may be due to the high levels of coverage achieved and a lower minimum threshold of alternative allele frequency (0.2%) utilised in this study. Interestingly, *Polg*^wt/wt^ mice displayed higher average alternative allele frequencies than *Polg*^D257A/D257A^. This effect, at first observation, is in stark contrast to what has been recorded previously and is not what one would expect from the literature. However, the high number of low-frequency heteroplasmies identified in *Polg*^D257A/D257A^ tissues may have caused a decrease in the overall average heteroplasmy levels (average alternative allele frequency) compared to *Polg*^wt/wt^ as a result of the high sequencing depth achieved across all samples. This result highlights the importance of looking at multiple measures of heteroplasmy to obtain an accurate assessment of the levels of mitochondrial DNA mutation that are present in a sample.

Mitochondrial DNA enriched *via* lrPCR amplification had a higher number of heteroplasmic sites and cumulative heteroplasmic burden than DNA enriched using the Mito-SiPE method. This difference was larger in *Polg*^D257A/D257A^ mice than in *Polg*^wt/wt^. There are two likely explanations for this observation. The first, that rounds of PCR amplification cause PCR errors that are subsequently identified as mitochondrial heteroplasmy, although a high-fidelity polymerase is used to negate this impact as much as possible. The second mechanism is that NUMT regions in the nuclear genome are being co-amplified and thus are mistaken for heteroplasmic, mitochondrial reads. If the first mechanism was the leading cause, it is unexpected that the difference would be higher in *Polg*^D257A/D257A^ tissues than in *Polg*^wt/wt^, as the error rate of the polymerase used for amplification should remain constant. Co-amplification of NUMT regions, however, could be affected by changes in mtDNA copy number – a feature that has previously been identified in *Polg* mutator mice^41–43^. Changes in mtDNA copy number may directly affect the amount of mispriming of NUMTs that occurs during lrPCR. Whilst our results cannot rule out either mechanism, this evidence suggests that co-amplification of NUMT regions as a more likely/influential mechanism for what would be interpreted as false positive heteroplasmic variants identified in PCR amplified samples. The metric of average alternative allele frequency did not display this same pattern but as explained previously, it is influenced by the presence of many apparent, low-frequency heteroplasmies that are identified when using lrPCR amplification at these sequencing depths and heteroplasmy threshold. This effect was even more dramatic when lrPCR was compared to mtDNA enrichment prep of C57BL6 wild-type tissues. This is largely due to the high baseline of mutation that is present in *Polg*^wt^ due to the intermediate phenotype of female breeder mice.

Finally, a tissue-specific effect on the levels of heteroplasmy was observed using the Mito-SiPE method in *Polg*^D257A/D257A^ mice. Spleen and colon samples had higher levels of heteroplasmy compared to heart, kidney, liver and lung, whereas, brain samples had lower levels. Heart, kidney, liver and lung were indistinguishable from one-another. This is an interesting finding, as previous studies have identified strong, tissue-specific effects of mitochondrial heteroplasmy in more tissues than what is reported here^36,44,45^. It is possible that enrichment methods used in previous studies have identified false heteroplasmic variants due to NUMT co-enrichment, and thus the differences observed are due to mtDNA copy number changes, rather than true mtDNA mutations. It is also possible, however, that the levels of coverage and low thresholds used in this study have led to this difference.

The limitation of this methodology is that it requires the availability of tissue for enrichment and as such it may not be feasible for archived DNA samples. DNA purified from intact cells and tissues using standard methods are predominantly nuclear DNA. These preparations and already existing NGS data are not compatible with this method, although such datasets are likely to have elevated NUMT contamination. However, with the explosion of interest in mtDNA and increasing levels of heteroplasmy-focused research, this method will provide an important tool for future explorations of mtDNA variation and its role in ageing and disease.

In conclusion, differential centrifugation and alkaline lysis may be used to enrich mitochondrial DNA eliminating the need forPCR amplification or probe hybridization. Avoiding sequence-dependent techniques greatly reduces the effect of NUMT contamination, a problem which has been identified in previous studies^23,24,28,30^. This technique, in addition to a modified bioinformatics pipeline, can be applied to different tissues and achieves ultradeep sequencing coverage. It provides a straightforward and robust workflow to assess heteroplasmy in mitochondrial DNA. This technique outperforms long-range PCR amplification and negates the potential impact of PCR errors and NUMT contamination on heteroplasmy analysis.

## Abbreviations

NUMT: Nuclear mitochondrial sequences
mtDNA: Mitochondrial DNA
POLG: Polymerase gamma

## Legends

**Supplementary figure 1.**
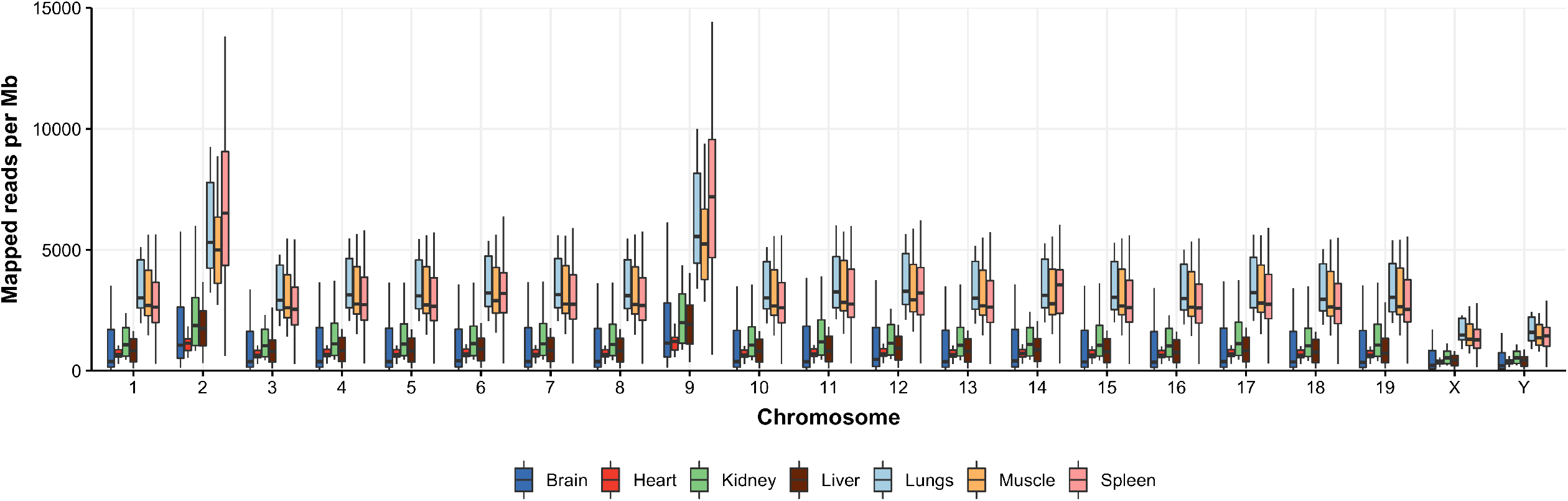
The distribution of nuclear contamination across the genome shown as boxplots for each tissue.

**Supplementary figure 2.**
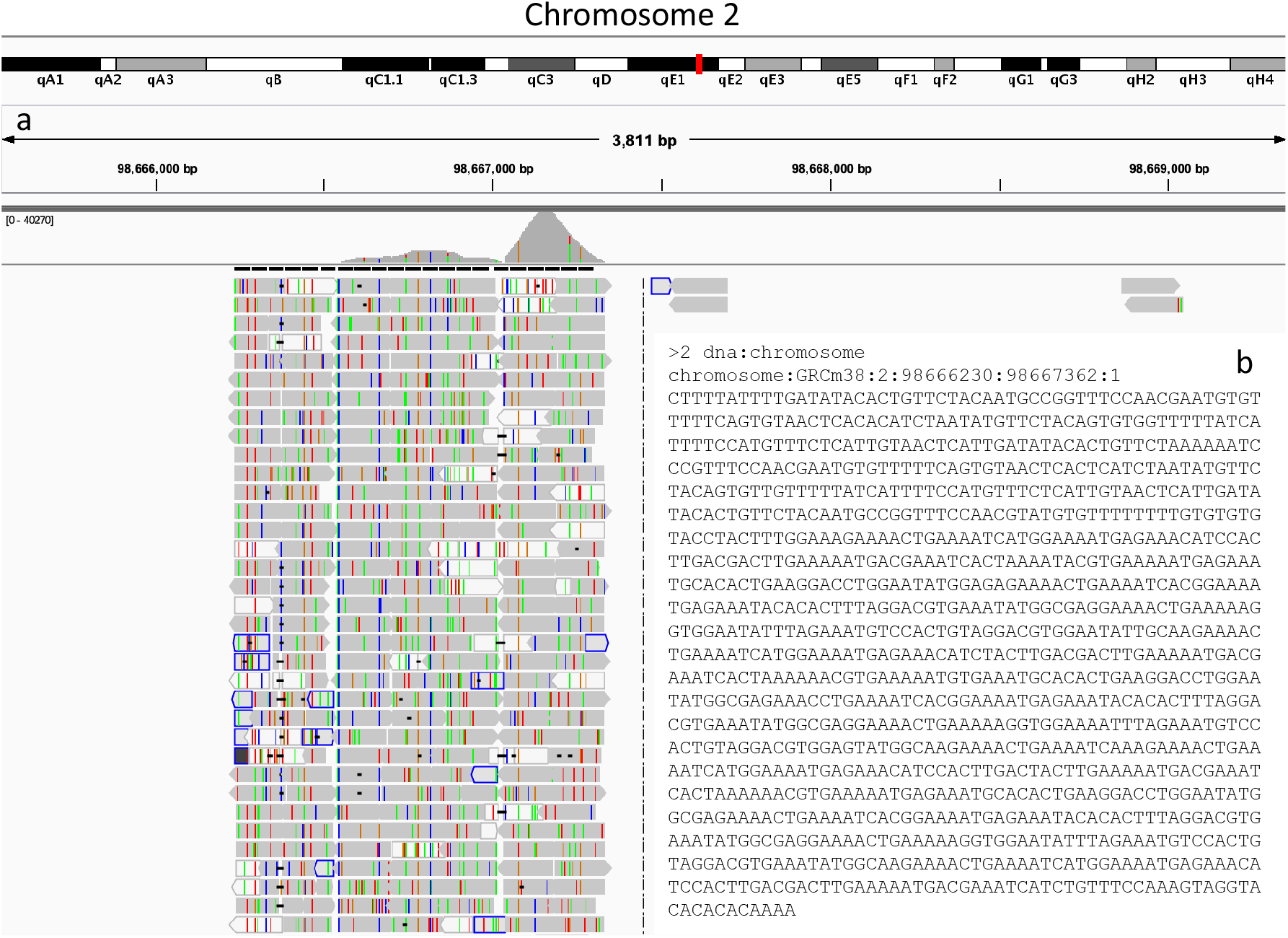
Sequence and alignment assessment of high coverage region on chromosome 2. **a**, Integrated genome viewer (IGV) image of high-coverage region on chromosome 2. **b**, The sequence of high coverage region in fasta format.

**Supplementary figure 3.**
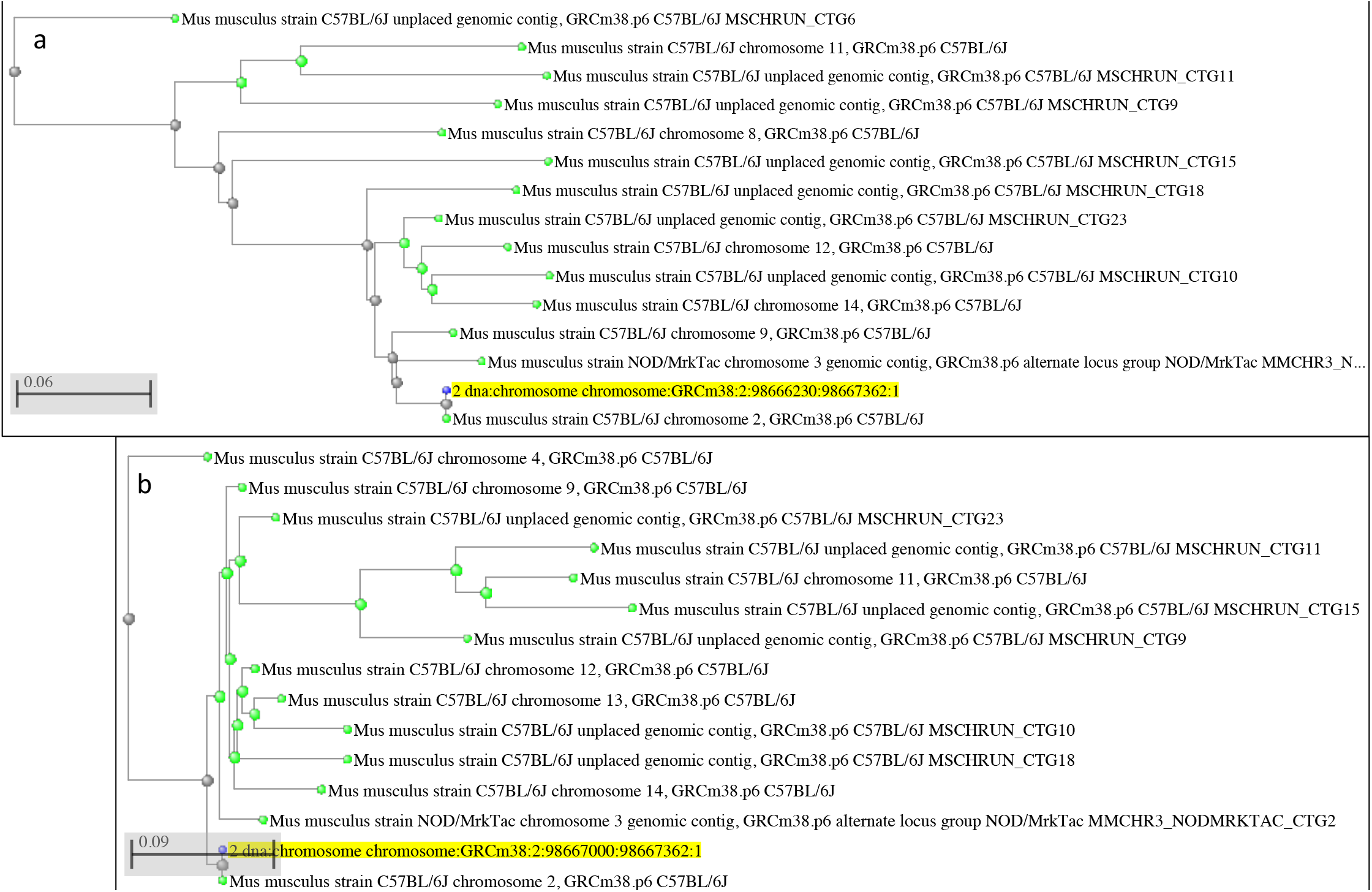
Phylogenetic tree of the blast results obtained from high-coverage sequence on chromosome 2. **a**, Phylogenetic tree of high coverage sequence chromosome 2:98666230-98667362. **b**, Phylogenetic tree of high coverage sequence chromosome 2:98667000-98667362.

**Supplementary figure 4.**
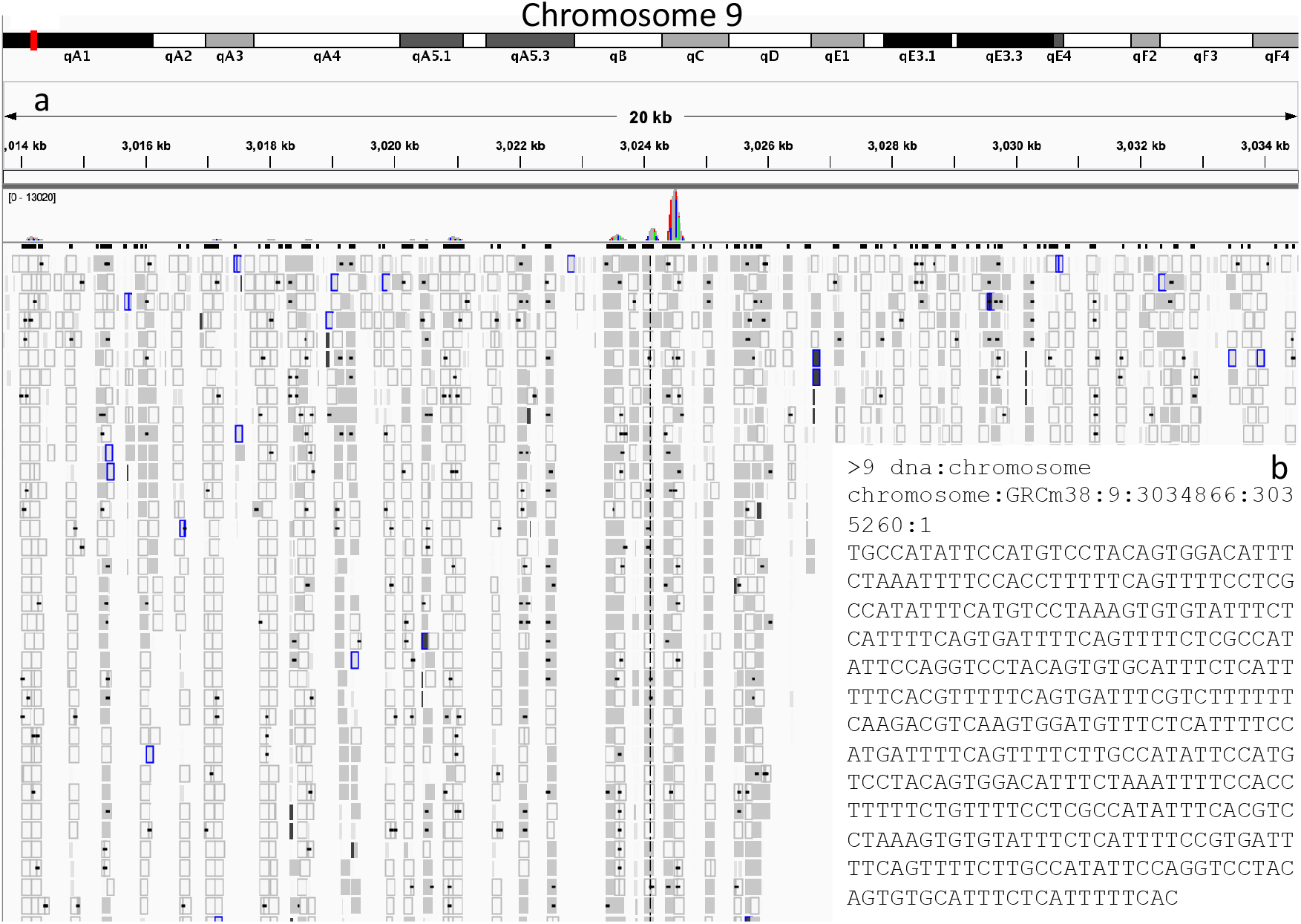
Sequence and alignment assessment of high coverage region on chromosome 9. **a**, Integrated genome viewer (IGV) image of high-coverage region on chromosome 9. **b**, The sequence of high coverage region in fasta format.

**Supplementary figure 5.**
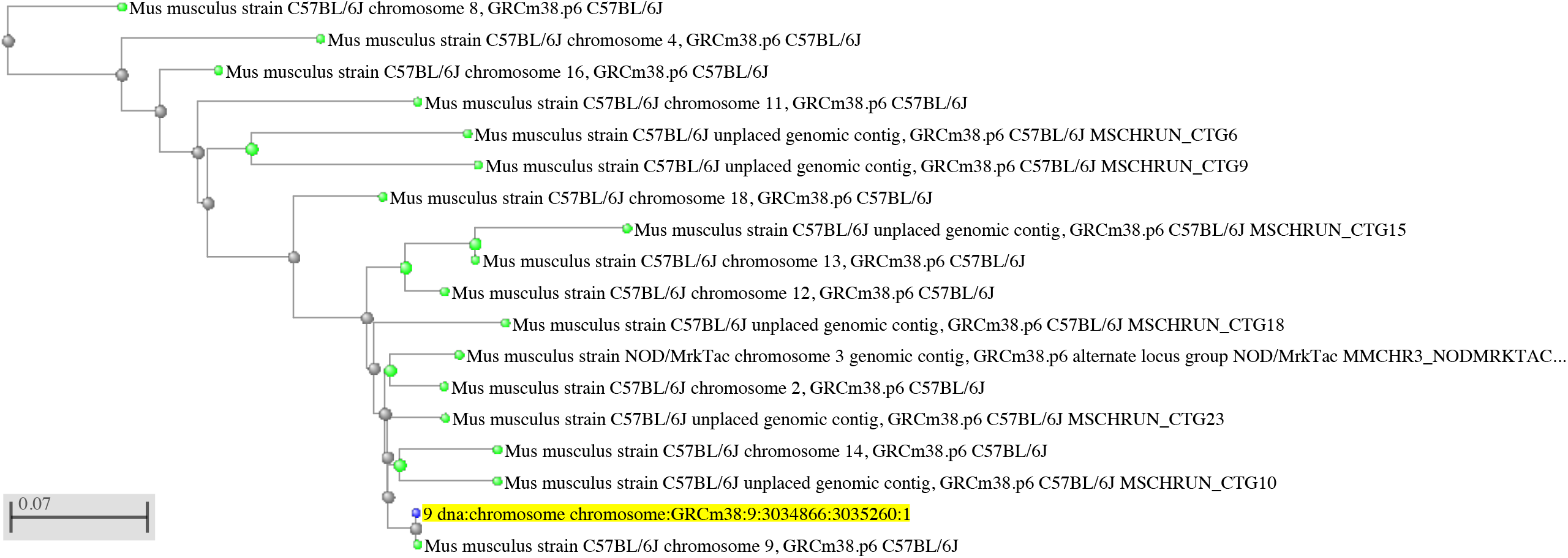
Phylogenetic tree of the blast results obtained from high-coverage sequence on chromosome 9. **a**, Phylogenetic tree of high coverage sequence chromosome 9:3034866-3035260.

**Supplementary figure 6.**
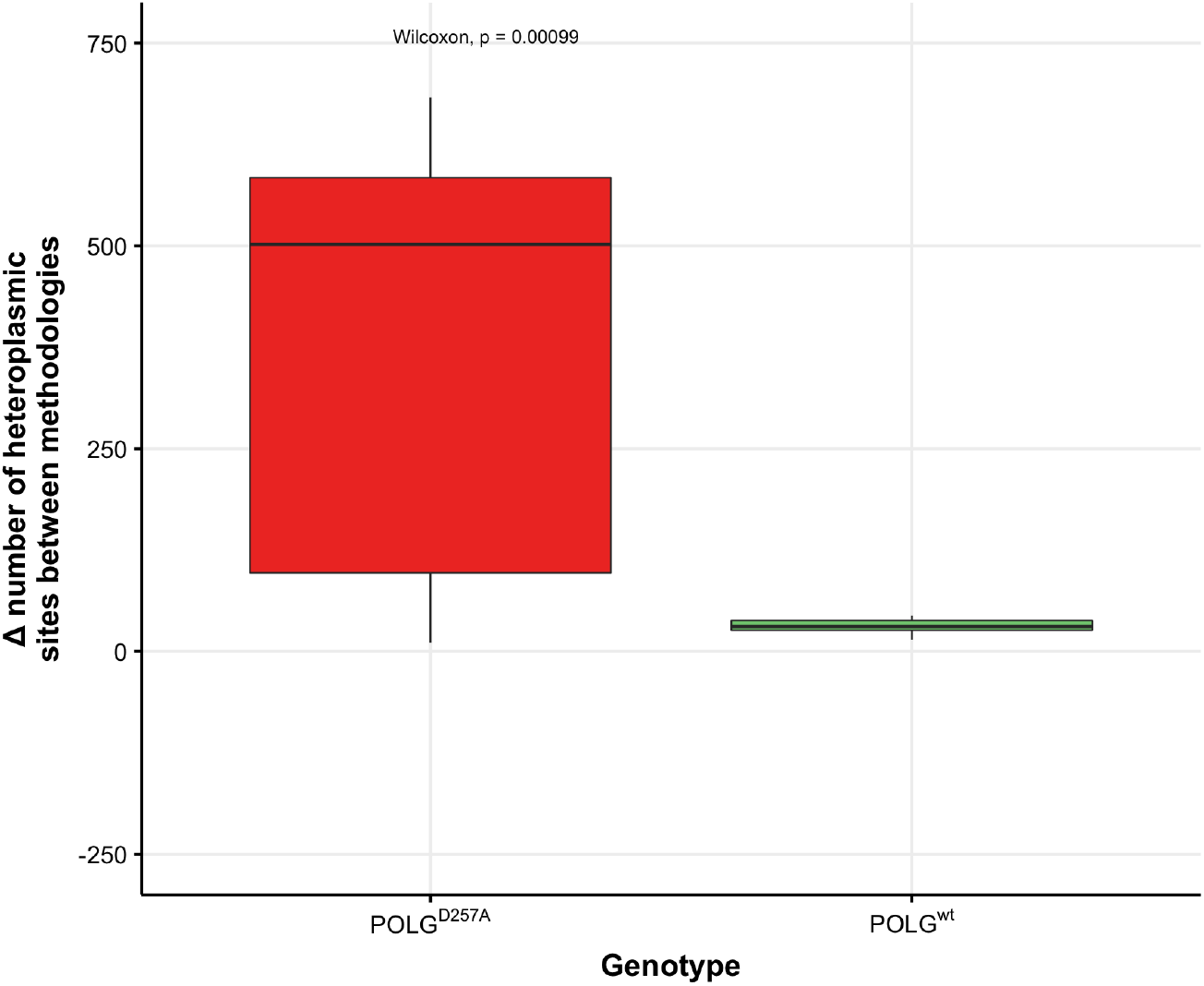
The difference in number of heteroplasmic sites found between lrPCR and mtDNA preparations in *Polg*^*D257A/D257A*^ and *Polg*^*wt/wt*^ tissues shown as boxplots.

## Methods

### Breeding and tissue harvesting

All animal protocols were reviewed and approved by the National Human Genome Research Institute (NHGRI) Animal Care and Use Committee prior to animal experiments. Mice were housed in shoe box cages and fed ProLab RMH 1800 diet (PMI Nutrition International) containing 50 μg vitamin B12/kg of diet and 3.3 mg folic acid/kg of diet. Breeding mice were fed Picolab Mouse Diet 20, containing 51 μg vitamin B12/kg diet and 2.9 mg folic acid/kg of diet. Heterozygous *Polg*^wt/D257A^ males and *Polg*^wt/D257A^ females were mated. Homozygous mutant and wild-type progeny were aged to 6 months at which point they were sacrificed. There were 4 *Polg*^wt/wt^ mice (2 male, 2 female) and 4 *Polg*^D257A/D257A^ mice (2 male, 2 female) used in the *Polg* experiments. Brain, heart, lung, liver, spleen, kidney and muscle tissue was isolated and mitochondrial DNA enrichment was performed on all tissues.

### Tissue homogenisation

Harvested tissue was placed in a homogenization tube with 10X volume per gram of fresh homogenization buffer i.e. 5 ml buffer for 500 mg tissue. Tissues were homogenised until no discernible whole tissue was present. The homogenate was then transferred to 1.5 ml microcentrifuge tubes and spun at 1000 g for 1 minute at 4°C. The supernatant was transferred to a new microcentrifuge tube and spun at 12000 g for 10 minutes at 4°C to pellet mitochondria. The mitochondrial pellet was resuspended with 100 µl of resuspension buffer for storage or for immediate DNA extraction.

### Mitochondrial DNA isolation

The mitochondria resuspension was added to 200 µl alkaline lysis buffer, vortexed, and placed on ice for 5 minutes. Potassium Acetate Buffer (150 µl) was then added and the mixture was vortexed slowly and placed on ice for 5 minutes. The mixture was centrifuged at 12,000 g for 5 minutes at 4°C to pellet proteins and the supernatant was decanted to a new tube. RNase (1 µg) was added to the mixture and left at room temperature for 15 minutes. Phenol-chloroform (500 µl) was added to each tube, inverted and placed on a shaker/rotator for 20 minutes. Afterwards, centrifugation at 12000 g for 2 minutes at room temperature was carried out. The aqueous (top) layer was decanted to a new tube (approx. 450 µl from this phase was retrieved) and 40 ul sodium acetate, 1 µl glycogen (20 mg/ml) and 1200 µl 100% EtOH were added. The mixture was inverted and mixed well then left on dry ice for 60 minutes. The mixture underwent centrifugation at 12,000 g and the supernatant was removed. The pellet was finally washed twice using 70% ethanol, air-dried, and resuspended in a low-TE buffer for sequencing or regular TE buffer for (q)PCR.

### Library preparation and next generation DNA sequencing

Libraries were generated from approximately 50 ng genomic DNA using the Accel-NGS 2S Plus DNA Library Kit (Swift Biosciences) using 5 cycles of PCR to minimize PCR bias. The DNA samples were sheared by sonication (Covaris Inc., Woburn, MA) to a mean of 300 bp. Libraries were tagged with unique dual index DNA barcodes to allow pooling of libraries and minimize the impact of barcode hopping. Libraries were pooled for sequencing on the NovaSeq 6000 (Illumina) to obtain at least 7.6 million 151-base read pairs per individual library. Sequencing data was processed using RTA version 3.4.4.

### Data processing and alignment

Fastq files were aligned to the mouse reference genome, GRCm38, using bwa mem using the default parameters^46^. Picard tools were used to add read groups, and to mark and remove duplicates^47^. Samtools was used to calculate the coverage across the nuclear and mitochondrial genome for each sample. Finally, R (v3.5.0) and ggplot2 (v3.3.0) were used for statistical analysis and subsequent visualisation of graphs^48,49^.

### Quantification of mtDNA copy number

Mitochondrial DNA copy number was assessed via qPCR targeting both mt-16S and nuclear-encoded hexokinase (HK), similarly as previously described^50^. Briefly, 2.5 µl LightCycler® 480 SYBR Green I Master (Roche, Molecular Systems, Inc, Germany), 2 µl of DNA (20 ng/µl) and 0.5 µl primer mix were added in triplicate to a 384-well plate and the reactions were carried out by the QuanStudio 6 Flex (Applied Biosystems, Foster City, CA, USA). The conditions were as follows: 95°C for 5 min, 45 cycles of 95°C for 10 s, 60°C for 10 s and 72°C for 20 s. A melting curve was performed using 95°C for 5 s, 66°C for 1 min and gradual increase to 97°C. Mitochondrial DNA copy number was assessed using the following formula: 2 × 2^ΔCt^ were ΔCt = Ct(mtDNA gene)−Ct(nDNA gene).

### Long-range PCR enrichment of mtDNA

This technique was used to amplify human and mouse mitochondrial DNA in two fragments from a whole DNA extract. DNA was quantified *via* Nanodrop (Methods 2.2.7) unless otherwise stated. Each PCR reaction consisted of Q5 High-fidelity Polymerase (0.02 U/μl), 5X Q5 reaction buffer (1X), 10mM dNTPs (300 μM), 5μM Forward and Reverse primers (0.25 μM, Table 2.3; human, Table 2.4; mouse). Template DNA (100 ng) was added to each reaction except for the no-template control (NTC) but an equivalent volume of molecular biology grade water was added instead. The temperature cycles were as follows: 1 × 30 s denature 98°C, 25 × 10 s denature 98°C, 30 s annealing 66°C, 4 minutes 30 s elongation 72°C, 1 × 10 minutes elongation 72°C on a thermocycler. Both fragments were quantified using qubit and mixed in equimolar ratios.

### Solutions

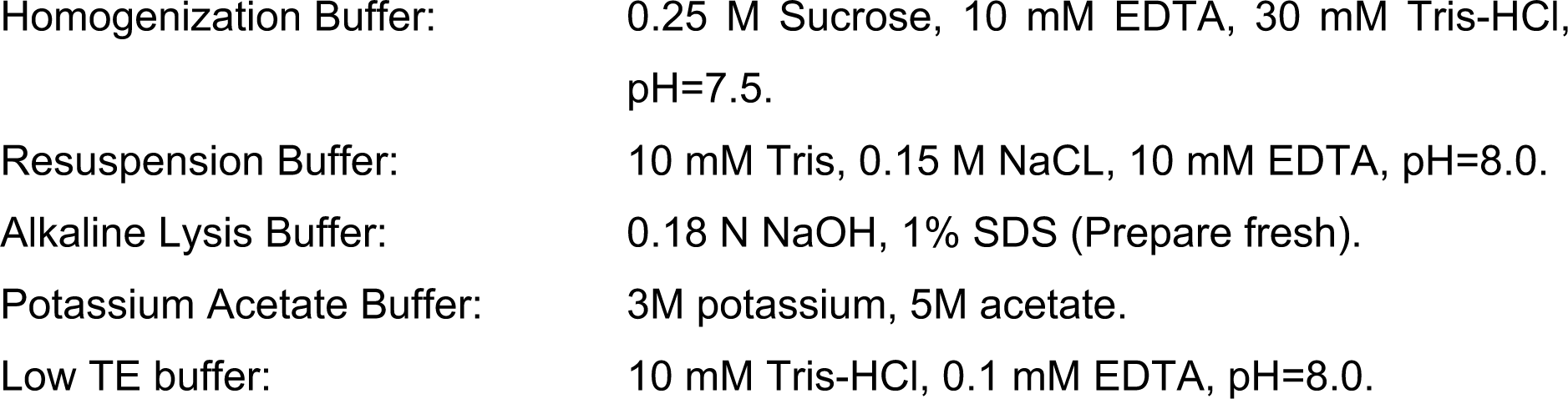

## Ethics approval and consent to participate

Not applicable

## Consent for publication

Not applicable

## Availability of data and materials

The sequencing datasets generated during and/or analysed during the current study are not publicly available due to their inclusion in a student PhD thesis but are available from the corresponding author on reasonable request. The datasets will be made public after submission of hard-bound thesis and subsequent graduation. The summary data, sequencing-depth statistics and heteroplasmy results were provided at time of submission.

## Author Contributions

DW, ME and DB performed the sample collections and enrichment. DW and DH performed bioinformatics and subsequent analysis. DW, DB, FP, DH, APMcD and LB contributed to the study design, data interpretation and prepared manuscript. All authors read and approved the final manuscript.

---

**S7.**
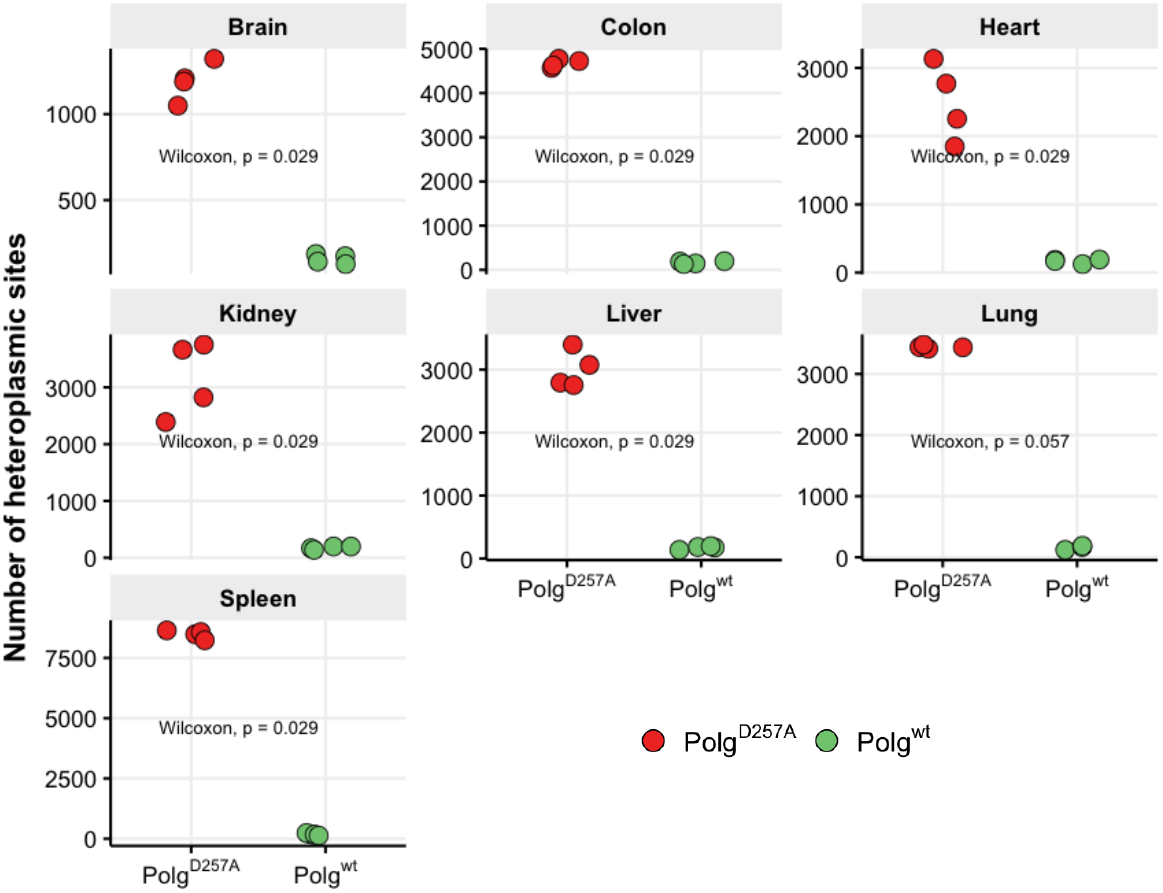

**S8.**
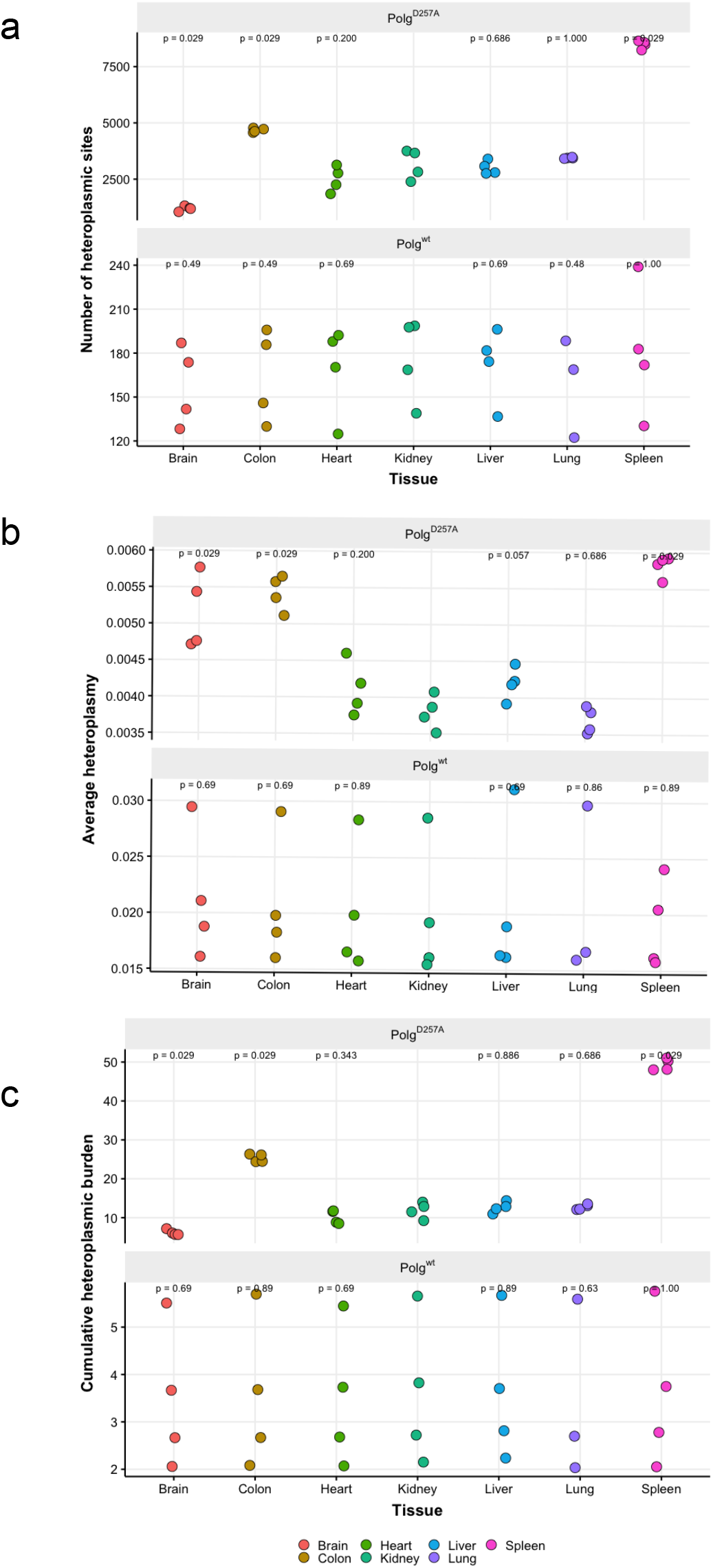

